# An Omics Analysis Search and Information System (OASIS) for Enabling Biological Discovery in the Old Order Amish

**DOI:** 10.1101/2021.05.02.442370

**Authors:** James A Perry, Brady J Gaynor, Braxton D Mitchell, Jeffrey R O’Connell

## Abstract

The “Omics Analysis Search and Information System” (OASIS), developed at the University of Maryland School of Medicine, enables discovery by allowing researchers to mine results from genome wide association studies (GWAS). When interesting signals are found, the research can immediately ask follow-up questions and get answers in real-time. OASIS provides this unique capability with a web-based, scientist-friendly search system and a variety of real-time analysis tools (linkage disequilibrium calculations, conditional analysis, and direct variant comparison) plus on-demand visualizations (boxplots, histograms, LocusZoom & Haploview plots, and pedigree charts). Because OASIS uses a web-based user interface, an understanding of programming or the UNIX operating system is not required. The OASIS application has been used to enable discovery from whole-exome, whole-genome, metabolome, transcriptome and methylome association results for Old Order Amish studies at the University of Maryland School of Medicine.

## INTRODUCTION

Millions, perhaps billions, of dollars are being spent on whole genome sequencing (WGS), whole exome sequencing (WES), methylation sequencing (Methyl-seq), and transcriptome sequencing (RNA-seq) to allow association analysis to be run with hundreds of phenotypes related to disease that are collected from cohort studies and electronic health records. When significant associations are detected, investigators typically want “all known annotation and functional information” for the marker/probe. Many annotation resources have been developed to meet this need including Annovar [1], The Ensembl Variant Effect Predictor (VEP) [2], SnpEff [3], FAVOR[4], and WSGA [5]. Resources for variant function include the ENCODE Project [6-9] and the NIH Roadmap Epigenomics Program [10-12], which have mapped epigenetic and regulatory regions throughout the human genome, plus the GTEx Project [13-15] which provides association data to understand the effect of variants on gene expression. These resources are a wonderful set of assets, but they can quickly create a massive data mining challenge for the researcher attempting to assemble, search and assimilate thousands of annotations and predictions for the variants with meaningful association signals.

At the University of Maryland School of Medicine, we experienced this data assimilation challenge first-hand for our studies of the Old Order Amish. Managing the data volume from chip-based genotyping was difficult but the volume from WES and WGS for 6000 subjects was simply daunting. To address the challenge, we developed the “Omics Analysis Search and Information System” (OASIS).

## METHODS

OASIS was designed to handle data from genotyping chips and WES for 6000 Amish subjects, producing 500,000 to 1 million variants, and WGS for 1100 Amish subjects that produced 10 million variants. From these genotype datasets, we generated association results for 1200 different Amish phenotypes. The genetic association results for our Amish studies were calculated using the MMAP [16] (Mixed Model Analysis for Pedigrees and Populations) software program. A schematic of the omics annotation and analysis pipelines is shown in Figure 1.

**Figure 1.**
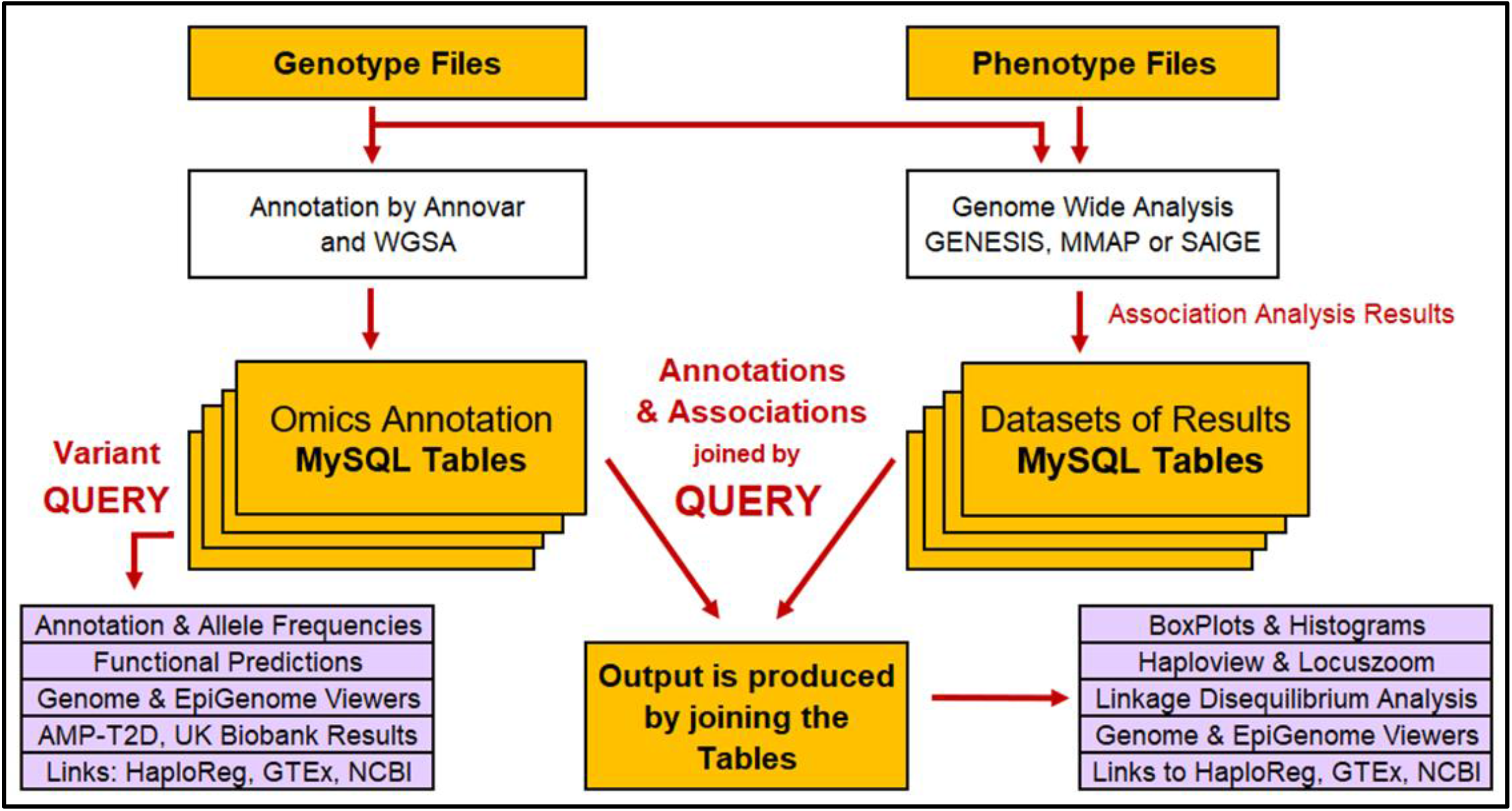
OASIS Annotation and Analysis Pipelines. OASIS files, tables and processes shown with gold backgrounds. Output from queries shown in purple. Processes external to OASIS shown in white.

Annotations for all genomic variants were collected from Annovar [1] and WGSA [5] and loaded into OASIS’ omics annotation tables of the MySQL database, shown on the left side of Figure 1. Next, association results from a genome wide association analysis were loaded into the MySQL tables shown on right side of Figure 1. The association analysis, which is performed external to OASIS, can be generated using any statistical analysis package. OASIS supports multiple file formats for association results including formats from Encore[17], GENESIS[18], and MMAP[16]. OASIS allows for two broad types of searches: 1) “query-by-variants” which searches just the functional annotations to help a user understand what variants are present in their study cohort (e.g. loss-of-function variants, variants reported as pathogenic by ClinVar) and the allele frequencies regardless of whether the study has phenotypes that associate with those variants, and 2) “query-by-trait” which provides a combined search of association results and functional annotations with criteria specified for both.

### OASIS Features

When performing either type of query, the OASIS user interacts with a web-based form to specify the search criteria plus the desired annotation and functional prediction information to be displayed. Upon form submission, OASIS issues a call to the MySQL database and, in the case of query-by-trait, performs a relational database join (see Figure 1) to generate the desired report. Along with the association results (e.g. p-value, effect size) and annotation for each variant, OASIS provides hyperlinks for each variant/gene to NCBI databases [19] (dbSNP, Gene, Variation Viewer), Broad (HaploReg [20], GTEx [14]), and Open Targets[21], and the International Mouse Phenotype Consortium [22]. Users can also perform ‘on the fly’ analyses of individual level data for immediate follow-up on questions or hypothesis generated when reviewing variant associations. These analyses, performed in real-time with system calls to MMAP, include options for direct variant comparison and conditional analysis. The conditional analysis combines multiple variants and phenotypic covariates, all specified by the user via an OASIS web form, into a customized model to determine if an association is driven by single or multiple variants. Also included are linkage disequilibrium (LD) calculations relative to a user-specified reference variant as well as buttons for exploring LD with LocusZoom [23] and Haploview [24] (Figures 2 & 3).

**Figure 2.**
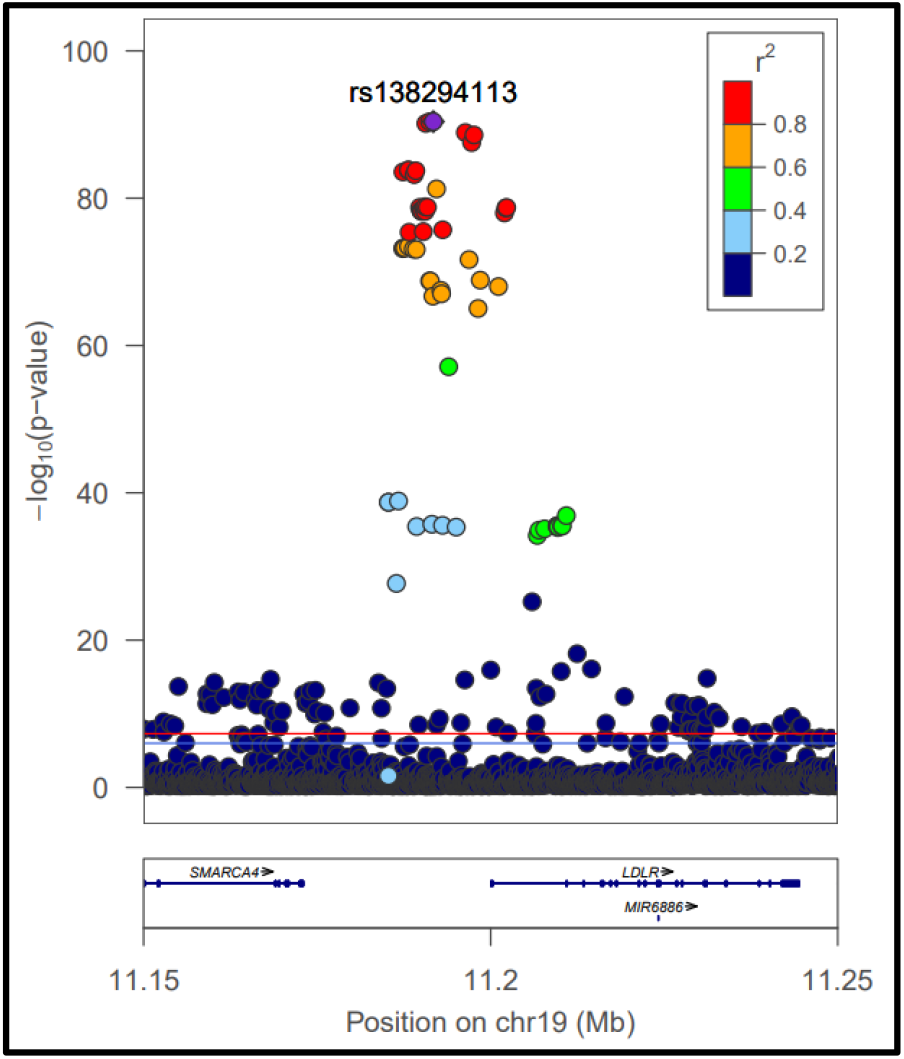
LocusZoom plot of LDL associations in promoter region upstream from *LDLR*.

Boxplots and histograms (Figure 3) showing trait values by genotype and split by sex are available with a single click. The boxplots are particularly useful as they also provide sample sizes for each genotype and variation within the genotype to allow the user to evaluate at a glance the robustness of the association result.

**Figure 3.**
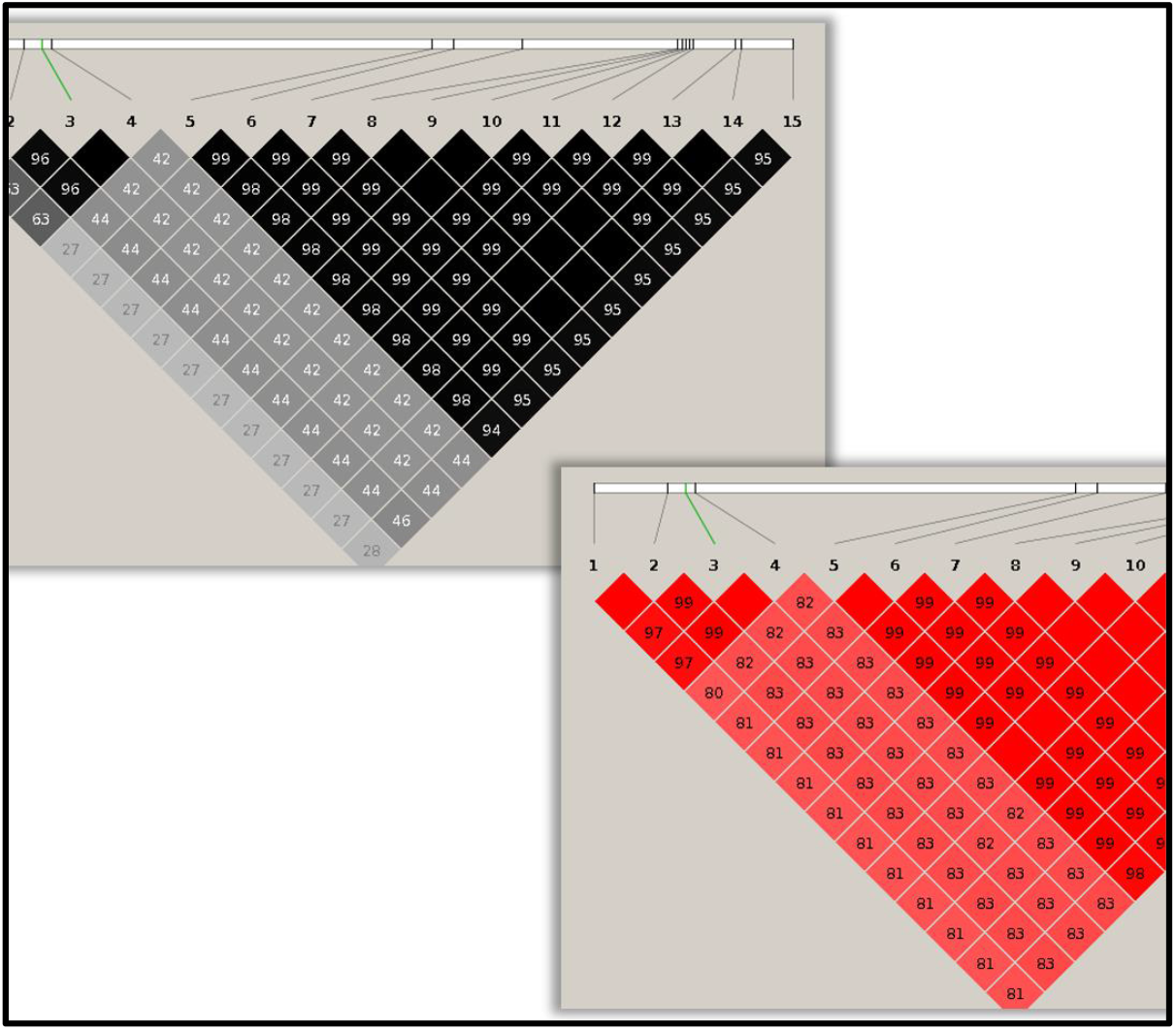
Haploview plot of linkage disequilibrium in Amish for region in *APOB*. r^2 (black), D-prime (red)

**Figure 3.**
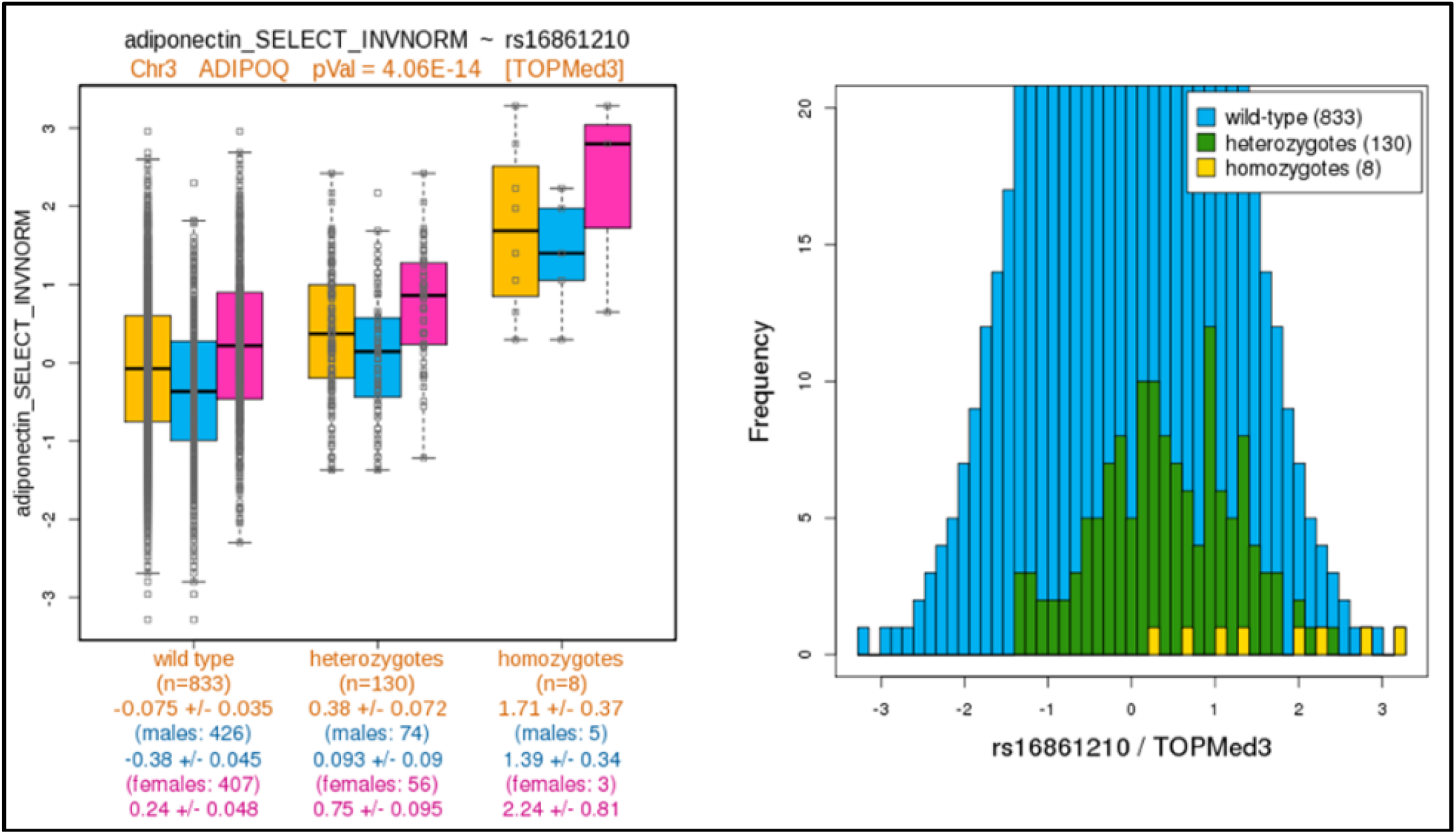
Boxplot and Histogram available from OASIS for any variant and phenotype combination. Boxplot on left is inverse normalized adiponectin by genotype for males (blue), females (pink) and both sexes (gold). Histogram on right shows frequency of phenotype values for each genotype including wild-type (blue), heterozygotes (green) and homozygotes (gold).

Genomic visualizations for multiple variants can be achieved by selecting desired variants in the OASIS report and then clicking the “UCSC” button. OASIS then sends the coordinates and color code information for each selected variant to the UCSC genome browser [25] along with the desired regulatory tracks for the tissues selected by the user.For example, in Figure 4, the user selected six variants that are upstream of the LDLR gene. One of the six lies directly in a DNase hypersensitivity site and also in a predicted enhancer region (yellow) for adult liver tissue, based on the chromatin HMM models from Roadmap [11]. This information can help with developing hypothesis on which variant is causal and what the functional mechanism might be.

**Figure 4.**
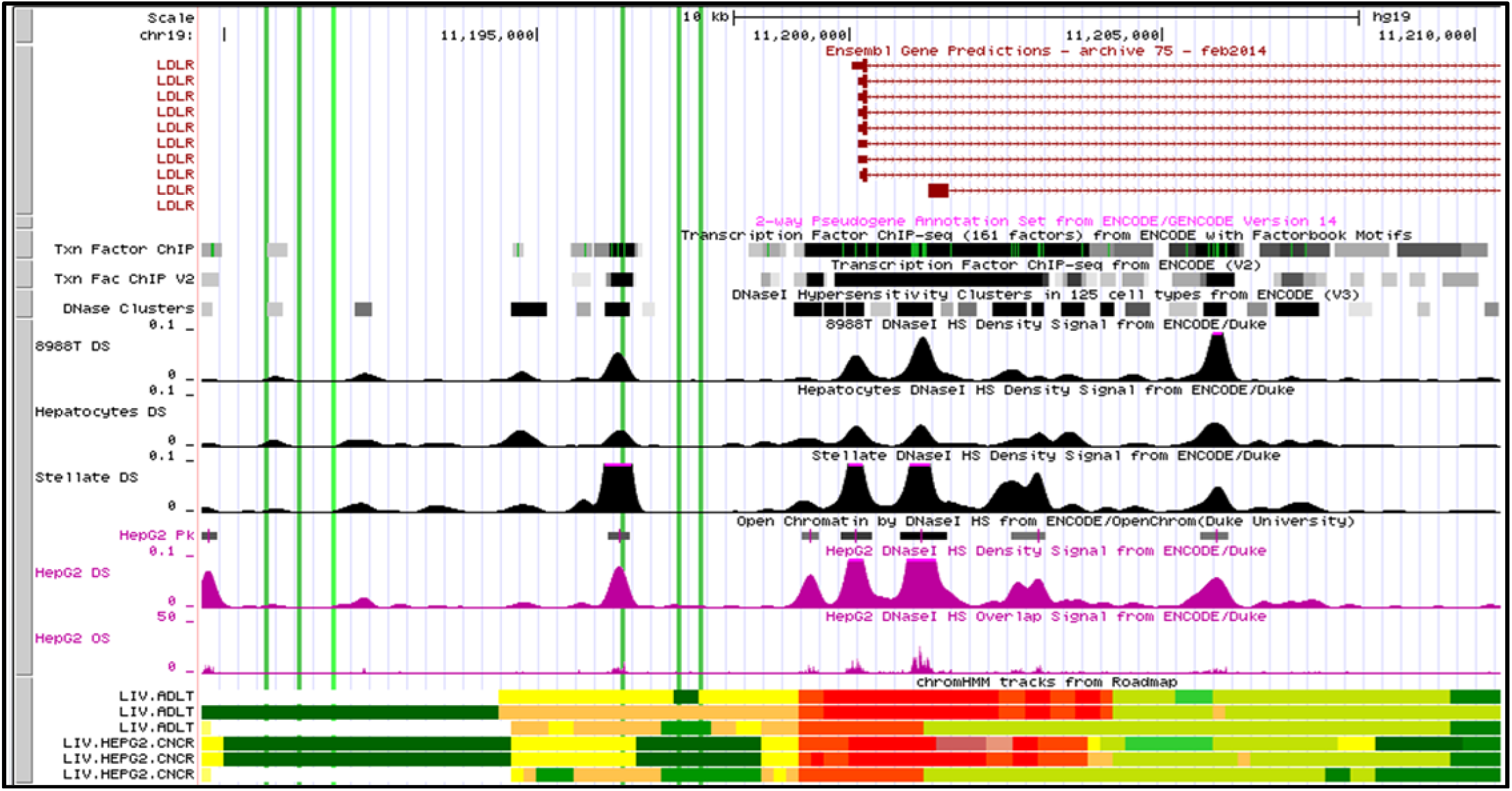
UCSC Genome Browser with user-specified variants (green lines) and tissue tracks (liver tissues)

Complete pedigree data is available for our Amish studies. This allows OASIS to generate pedigree charts by genotype for any variant and include the numeric phenotype values for each subject as shown in Figure 5. The pedigree chart feature is available with one additional click after drawing a boxplot. This feature can be used for any cohort with a pedigree data file.

**Figure 5.**
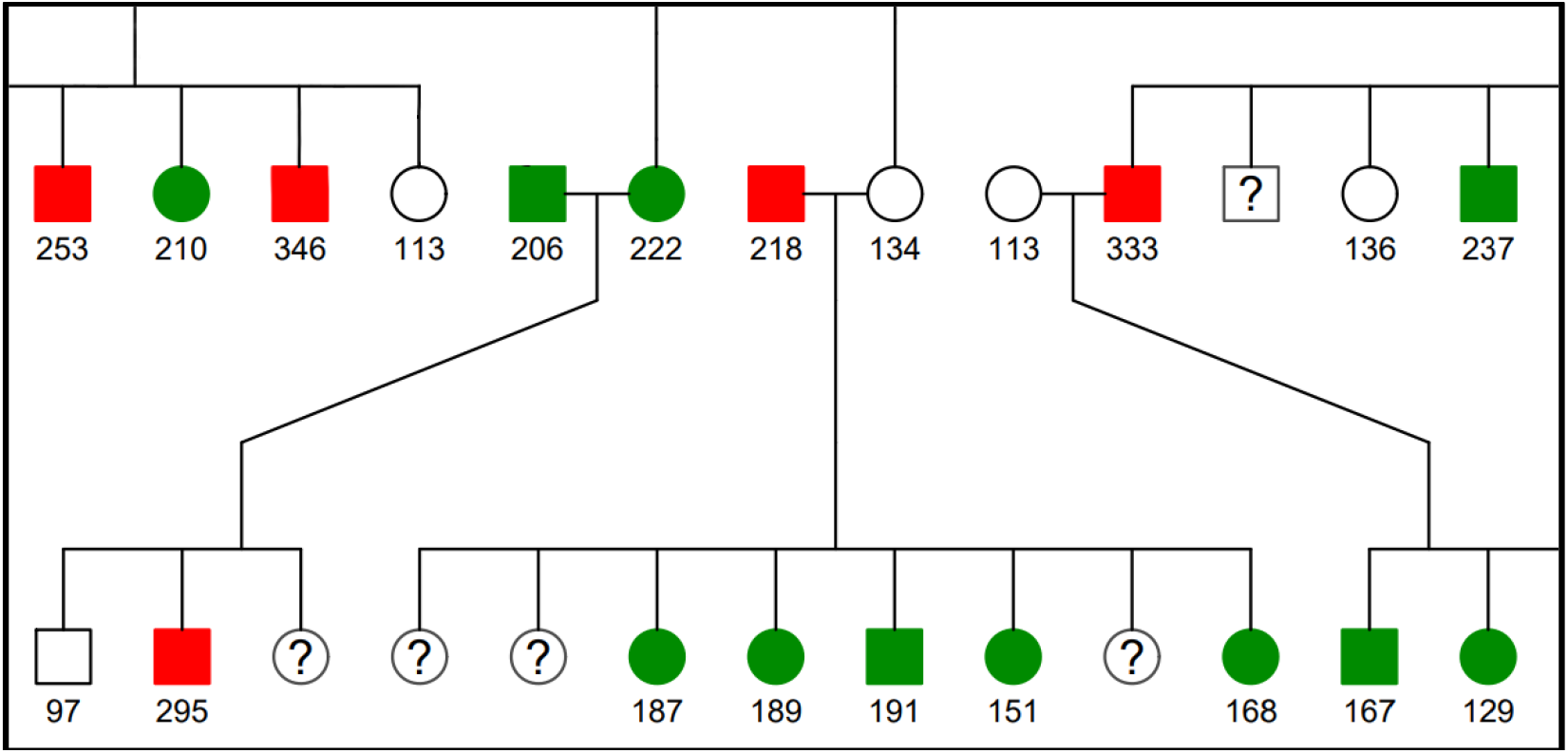
OASIS generated pedigree chart for a selected variant showing subjects by genotype. Homozygous-reference (white), heterozygous (green), homozygous-alternate (red), unknown genotype (question mark), male subjects (squares), female subjects (circles), connecting lines indicate parent-child-sibling relationships. Numeric values are subject measurements for the selected phenotype.

### Under the hood

OASIS is a web-based application that runs on a Linux/Apache [26] web server with 12 dual-core, 128Gb memory, and 8TB disk storage. The user interface is form-based and is displayed via a web browser using HTML5, CSS3 and JavaScript [27]. Visualizations (boxplots, bar charts, pedigree charts) are written in R [28]. The core code is written in Perl [29] using Perl libraries CGI, DPI and TMPL for preparing and processing requests and responses to/from the Apache web server and a MySQL [30] database. The Perl code also generates the customized links to online resources (NCBI, GTEx, etc.) and, if the “UCSC button” is clicked, will integrate user-selected variants into customized calls to the UCSC genome browser [25] to provide an understanding of how the variants are related to other genomic features with tissue-specific tracks in the UCSC genome browser. The tracks displayed in the UCSC genome browser are specified through OASIS’s own customized setup files for a dozen tissues and/or diseases (e.g. liver, heart, diabetes) for the user to choose. Open source packages, such as LocusZoom [23] and Haploview [24], are installed on the web server and called by the core code using selections and options provided by the user via web page forms.

Data storage for OASIS is maintained in multiple formats for efficiency. These include MySQL database tables, standard comma-separated-value (csv) files and compressed binary genotype/omics files. MySQL database tables, with multiple indexes, are used to store the annotated omic variants/probes and the association results. Current prototypes have omics annotation tables with as many as 468 million variants. The association results tables contain the variant/probe identifier, used as the join field for MySQL queries joining to the omics annotation table, plus results fields from the association analysis. Association results tables are split by study for efficiency.

Standard csv files are used to store subject-level phenotype data needed for the real-time generation of boxplots, bar charts and LD calculations, all based on the limited set of subjects for any given phenotype. Phenotype descriptions, sample annotation files (with age, sex, ancestry, etc. for each sample), and pedigrees for the Amish studies are also stored in standard csv files. Genotype/omics files, also needed for boxplots and LD calculations, are converted to highly compressed binary formats created by MMAP. Genotype files for WGS are divided by chromosome for efficiency.

## RESULTS and DISCUSSION

OASIS for our Amish cohort currently contains 2 billion “searchable” association results generated from 27 million genotypes. Phenotypes include 1200 traits from the Amish research studies, 200 traits from metabolomics datasets, methylation M-values for 450,000 probes and expression levels for 15,000 RNA-Seq transcripts. Phenotypes may be continuous (LDL, BMI) or binary traits such as “diabetes status”. In the case of eQTL or mQTL analysis, the expression or methylation level becomes the trait. Metabolomic, eQTL and mQTL association results have also been successfully implemented in OASIS.

OASIS is widely used at the University of Maryland by the Amish Research team and their collaborators. During a recent 12-month, 35 Amish researchers performed 30,000 separate OASIS queries. All visualizations and real-time analyses are initiated from the web-based interface at the direction of the investigator who is not required to understand programming, command-line syntax or the Unix operating system. Visualizations available with a single click include boxplots, histograms, pedigree charts, Locuszoom and Haploview plots and UCSC Genome Browser displays for selected variants. Real-time analyses include conditional analysis between selected variants, LD calculations based on the cohort genotypes, and one-to-one comparisons for selected variants.

## SOFTWARE AVAILABILITY

We are currently seeking funding to produce a production version that includes comprehensive web security. When upgraded, OASIS will be available for licensing free of charge for non-commercial use. Please visit the following link for updates on OASIS software availability. http://edn.som.umaryland.edu/OASIS/

## ACKNOWLEDGEMENTS

This work was supported by the National Institutes of Health [U01HL137181].

